# Age-related alterations in functional and structural networks in the brain in macaque monkeys

**DOI:** 10.1101/2024.09.15.612815

**Authors:** Kazuya Ouchi, Shinya Yamamoto, Makoto Obara, Yasuko Sugase-Miyamoto, Tomokazu Tsurugizawa

**Affiliations:** Human Informatics and Interaction Research Institute, National Institute of Advanced Industrial Science and Technology, 1-1-1 Higashi, Tsukuba-City, Ibaraki 305-8568, Japan; Faculty of Engineering, Information and Systems, University of Tsukuba, Tsukuba, 305-8573, Japan; Graduate School of Comprehensive Human Sciences, University of Tsukuba, 1-1-1, Tennodai, Tsukuba, Ibaraki 305-8577, Japan; Philips Japan, 2-13-37 Kohnan, Minato-ku, 108-8507, Tokyo, Japan

**Keywords:** macaque monkey, functional connectivity, structural connectivity, network, MRI

## Abstract

Resting-state networks (RSNs) have been used as biomarkers of brain diseases and cognitive performance. However, age-related changes in the RSNs of macaques, a representative animal model, are still not fully understood. In this study, we measured the RSNs in macaques aged 3–20 years and investigated the age-related changes from both functional and structural perspectives. The proportion of structural connectivity in the RSNs significantly decreased, whereas functional connectivity showed an increasing trend with age. Additionally, the amplitude of low-frequency fluctuations tended to increase with age, indicating that resting-state neural activity may be more active in the RSNs may increase with age. These results indicate that structural and functional alterations in typical RSNs are age-dependent and can be a marker of aging in the brain.

## 1 Introduction

Resting-state networks (RSNs) in the brain that reflect cognitive function are dependent on aging in humans (Goelman, et al. 2024). Further, research has shown that macaque monkeys have RSNs homologous to those in humans (Hutchison, et al. 2011). Indeed, the previous study investigating age-related differences in RSNs among macaques, ranging in age from 1 to 15 years, reported that the RSNs of monkeys were not widely changed, but were fine-tuned (Rao, et al. 2021). However, this study did not investigate older monkeys. Another study that surveyed structural T1-weighted MRI scans of 66 macaques (aged 5–31 years) reported enlargement of the lateral ventricles and smaller volumes in the frontal cortex, caudate, putamen, hypothalamus, and thalamus in older macaques compared to their younger counterparts (Dash, et al. 2023). The previous study demonstrated that the regional cerebral blood flow and the regional metabolic rate of glucose in aged macaques were lower than those in young macaques (Noda, et al. 2002). The metabolic rate of glucose in specific brain regions, such as the hippocampus and prefrontal cortex, is associated with cognitive performance (Eberling, et al. 1997; Upright and Baxter 2021), and changes in brain metabolism with age are important factors. As neuronal and astrocyte activity require energy metabolism, altered energy metabolism should be relevant to the blood oxygenation level-dependent (BOLD) signal, a signal source of fMRI regulated by neuronal and astrocyte activity. The normal aging process in humans is accompanied by a restructuring of the functional connectivity (FC) networks, characterized by a shift from more segregated to more integrated brain networks, which has been found to be important for changes in our cognitive performance. In addition to the alterations in FC with age, human MRI studies have revealed age-dependent alterations in structural connectivity (SC) (Stumme, et al. 2022). However, the relationships among SC, FC, and age in macaques have not yet been fully investigated.

In addition to FC and SC, some indices were calculated from the voluntary fluctuations of functional magnetic resonance imaging (fMRI) signal changes depending on age. The amplitude of low-frequency fluctuations (ALFF), global connectivity, and local connectivity measured during RSN have also been associated with age-dependent metabolism in humans (Bernier, et al. 2017; Tomasi, et al. 2013). The ALFF is an index calculated from the square root of the power spectrum in the frequency range, generally between 0.008 and 0.1 Hz, and it has been validated as a useful biomarker of the brain’s physiological state (Yang, et al. 2007). Global correlation analyses have indicated correlations between the signal fluctuation at each voxel and that averaged in the whole brain, whereas local correlation has indicated the synchronization between a specific voxel and neighboring voxels. These have been utilized as indices to demonstrate the coordination between neurons (Deshpande, et al. 2009). However, whether these indices change with age in macaques remains unclear. We therefore hypothesized that brain function and structure changed depending on age in macaques and investigated age-related changes in SC, FC, ALFF, global connectivity and local connectivity within the RSNs in macaques.

## 2 Materials and Methods

### 2.1 Subjects

This study used seven macaques (Macaca fuscata, 4male and 3 female, age 3 to 20 years old). All experiments were approved by the Ethics Committee of the National Institute of Advanced Industrial Science and Technology.

### 2.2 MRI acquisition

All MRI experiments were performed using a Philips Ingenia 3T MRI system with an 8-channel brain array RF coil. Macaques were anesthetized using a combination of medetomidine (0.05 mg/kg), midazolam (0.3 mg/kg), and ketamine (0.4 mg/kg) during MRI scanning, which was injected intramuscularly 30 minutes before the start of the survey scan. Diffusion-weighted imaging (DWI) was performed using a diffusion-weighted spin-echo echo planner imaging (EPI) sequence with the following parameters: repetition time (TR)/echo time (TE) = 20,000/93 ms, voxel size = 1.2 × 1.2 × 1.2 mm^3^ /voxel, matrix size = 112 × 112, 44 slices, b-value = 0, 1,000, and 3,000 s/mm^2^, and 64 directions. Resting-state functional MRI images were acquired using a gradient-echo EPI sequence with the following parameters: TR/TE = 2,000/30 ms, flip angle = 80 °, voxel size = 1.3 × 1.3 × 1.3 mm^3^ / voxel, matrix size = 80 × 80, 35 slices, and a total of 10 min (300 volumes). The T1-weighted image was acquired for the registration in image in preprocessing, using a magnetization-prepared rapid gradient echo (MPRAGE) sequence with the following parameters: TR/TE = 13/5.8 ms, voxel size = 0.6 × 0.6 × 0.6 mm^3^/voxel, matrix size = 200 × 250 × 167.

### 2.3 Structural connectivity (SC)

Preprocessing of the DWI data, including denoising, distortion correction, eddy current correction, motion correction, and B1 field inhomogeneity correction, was performed using MRtrix3 (https://www.mrtrix.org/). Non-brain tissues were removed using FSL software (https://fsl.fmrib.ox.ac.uk/fsl/). Registration was conducted using the ANTS software (www.nitrc.org/projects/ants/). The National Institute of Mental Health (NIMH) (https://afni.nimh.nih.gov/) and Cortical Hierarchy Regions of Interest (ROIs) of Resus Macaques (CHARM) (https://afni.nimh.nih.gov/) were used. Fiber tracking was performed for each macaque using MRtrix3. We set the number of tracts in the entire brain at 100 million. A matrix of each subject’s connectome was created to extract the fiber connectivity between each brain region.

### 2.4 Functional connectivity (FC)

Statistical parametric mapping SPM12 software (Welcome Trust Center for Neuroimaging, UK) was used to preprocess the fMRI data, which involved slice timing correction, motion correction by realignment, normalization, and smoothing. The processed data were then processed using CONN toolbox (Whitfield-Gabrieli and Nieto-Castanon 2012). Signals related to the six affine parameters of head motion and signals within the white matter and cerebrospinal fluid were regressed-out. The residue of the fMRI signals was then detrended, and slow periodic fluctuations were extracted using a bandpass filter (0.008–0.09 Hz). The same ROIs as in the SC analysis were used to calculate the FC. Each component of the FC matrix was displayed as a Fisher-transformed bivariate correlation coefficient between the BOLD time series of a pair of brain regions.

### 2.5 Individual similarity of SC and FC

Similarities among participants were calculated using a homemade program in MATLAB. All combinations of SC and FC were transformed into one-dimensional vectors for each participant (Ouchi, et al. 2024). Pearson’s correlation coefficients for SC and FC were calculated as the similarity between pairs across all individuals (Hyon, et al. 2020).

### 2.6 Network analysis

We extracted the RSNs identified in a previous study using independent component analysis (ICA) (Rao, et al. 2021). The default mode network (DMN) is primarily located in the posterior medial cortex, superior parietal lobule, parieto-occipital area, and visual area. The sensorimotor network (SMN) is a local network of the sensorimotor cortices. The salience network (SN) is a connectivity in the anterior cingulate cortex, prefrontal cortex and caudate nuclei. The visuospatial network (VSN) mainly encompasses the parieto-occipital area and visual cortices. We used the FMRIB Software Library (FSL) Multivariate Exploratory Linear Optimized Decomposition into Independent Components (MELODIC) to perform probabilistic group ICA (Smith, et al. 2004). The multi-session temporal ICA concatenation approach was applied for all macaques in a temporally concatenated manner for ICA analysis. A total of 70 independent components were extracted from all macaques, and RSNs were extracted using the criteria described above. The RSNs obtained from ICA were then used as ROIs for the calculation of FC, SC, ALFF, global correlation, and local correlation in all individuals using the CONN toolbox (Whitfield-Gabrieli and Nieto-Castanon 2012). FC was calculated as the Pearson’s correlation coefficient between the time course of fMRI signals within the RSNs. The ALFF was further calculated as the square root of the power spectrum in the frequency range of 0.008-0.09 Hz within each RSN (Yang, et al. 2007). Global and local correlations involve calculating the average of correlation coefficients for each voxel in the brain. Global correlation represents the correlation of the averaged time course in the RSN with all other voxels (Saad, et al. 2013), while local correlation focuses on the correlation of the averaged time course in the RSN with voxels neighboring the RSN (Deshpande, et al. 2009).

### 2.7 Statistics analysis

The Pearson’s correlation coefficients between age and SC, FC, ALFF, and global and local correlations were calculated, and the statistical significance (p < 0.05) of the correlation was computed using regression analysis in MATLAB.

## 3 Results

### 3.1 Individual similarity of SC and FC

The averaged SC and FC of five young and two old macaques are shown in Fig 1. Statistical comparisons were not performed, because the sample size of the old macaques was too small (n = 2). The interhemispheric connections of the SC appeared to be reduced in the old group (Fig 1A). In particular, the connections between the left and right frontal and parietal lobes and the connection with some regions of the temporal lobe seemed to be reduced in the old group. In contrast, the FC was similar between the young and old groups (Fig 1B). We subsequently assessed the inter-individual similarity (Fig 1C). SC indicated high similarity within young macaques (r = 0.91-0.94), while old macaques show low similarity with young macaques (r = 0.77-0.82). The similarity of the FC was low among all subject, but with higher similarity between older macaques (r = 0.35) than others (r = 0.11-0.25).

**Fig 1.**
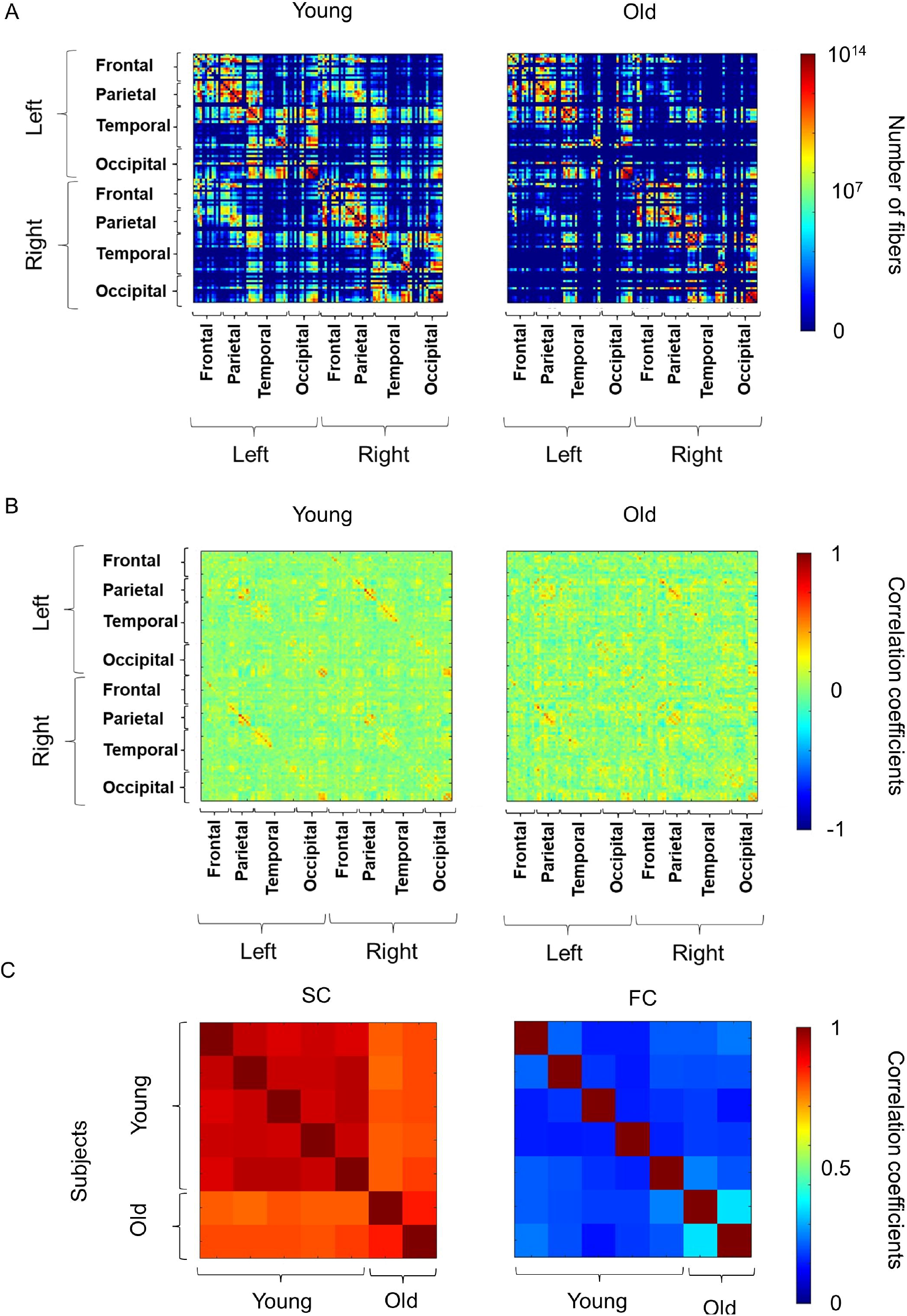
Overview of the connectivity matrix and similarity matrix. **(A)** The averaged structural connectivity (SC) in young and old macaques. The color bar represents the number of bundles between the regions of interest and “e” indicates Napier’s constant. **(B)** The averaged functional connectivity (FC) in young and old macaques. The color bar represents the value of Peason’s correlation coefficients between the regions of interest. **(C)** Individual variations of SC and FC in macaques and humans.

### 3.2 Age related changes in SC and FC

We analyzed the correlation between the functional and structural parameters of typical RSNs and age. Typical RSNs were successfully extracted from all macaques (Fig 2A). The proportions of SC and the average FC were calculated using these RSNs. The SC among DMN, SMN, and VSN showed higher proportion to SC in whole brain (Fig. 2B). The FC between RSNs showed high connectivity with VSN (Fig. 2C). Remarkably, significant negative correlations of SCs with age (r ≤ -0.8) were found in DMN and SMN (p = 0.025 and 0.015, respectively) (Fig 3A). Although the correlation between SC and age in the VSN was not statistically significant (p = 0.085 > 0.05), a negative correlation (r = -0.69) was nevertheless observed. In contrast, no correlation (r = 0.01) was observed between the SC and age in the SN. In contrast to the negative correlation between SC and age, FC was positively correlated with all RSNs (Fig 3B). The FC within the SMN showed a significant positive correlation with age (r = 0.77, p = 0.042) (Fig 3B). A positive correlation between age and FC in the DMN, SN, and VSN was also observed; however, the relationship was not significant (r = 0.35, 0.39, and 0.59, p = 0.45, 0.39, and 0.17, respectively).

**Fig 2.**
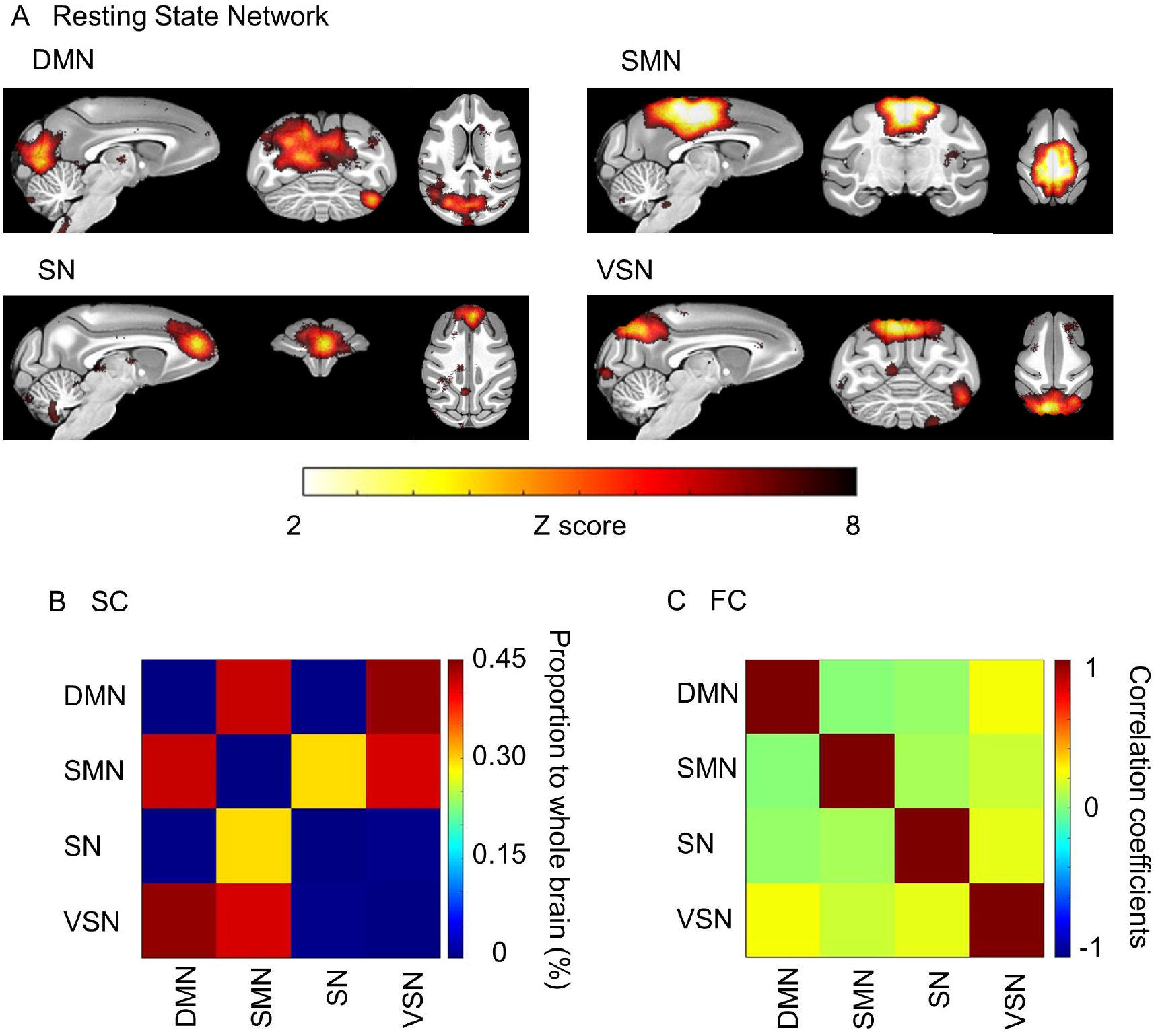
Resting state networks (RSN) extracted by independent component analysis. **(A)** ROIs of the Default mode, Sensorimotor, Salience, and Visuospatial networks were extracted. The color bar represents the value of the Z score. **(B)** The averaged structural connectivity matrix. The color bar represents the percentage of SC to the whole brain. **(C)** The averaged functional connectivity matrix between RSNs. The color bar represents Peason’s correlation coefficient.

**Fig 3.**
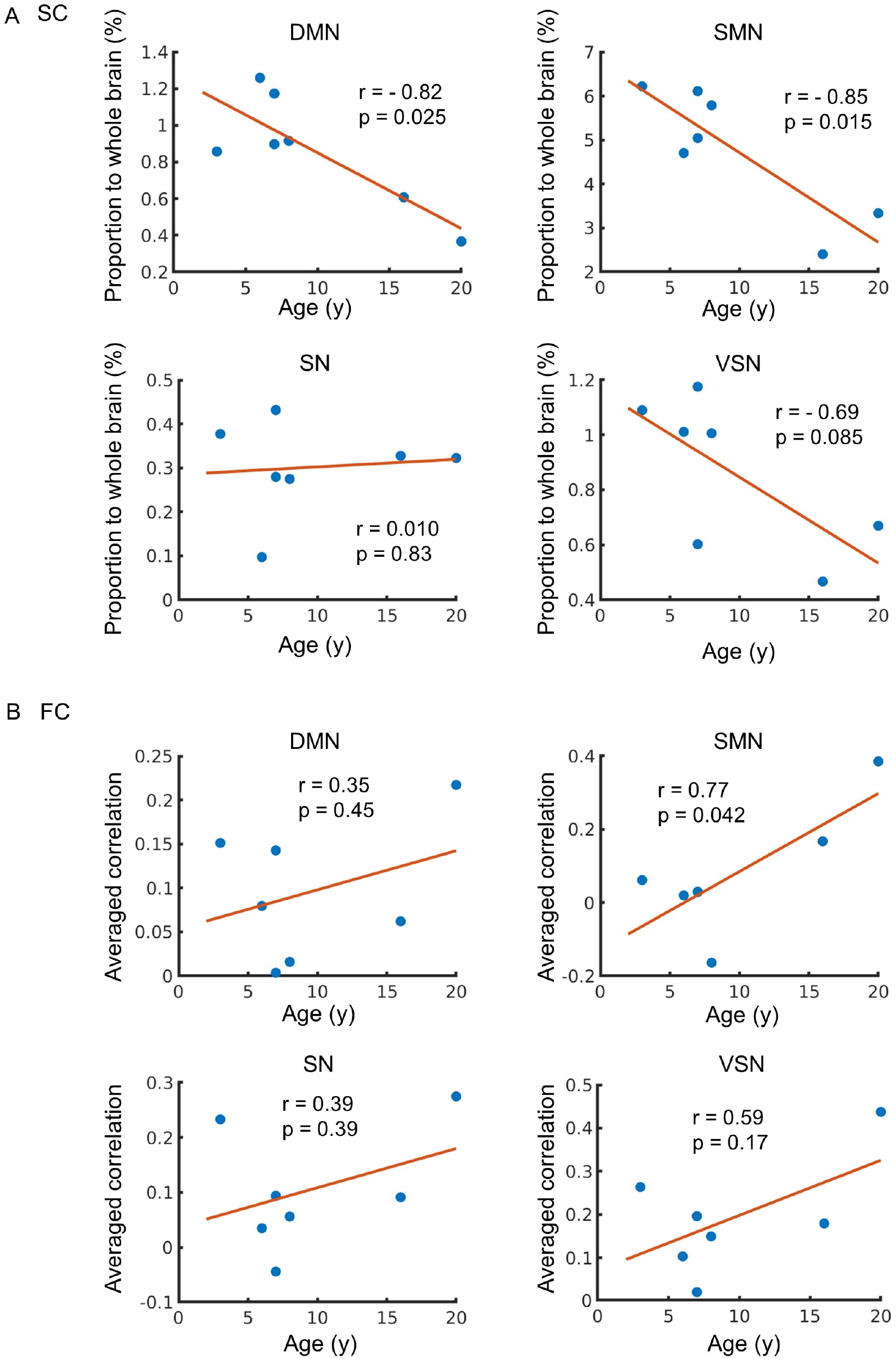
Age-related changes in structural connectivity (SC) and functional connectivity (FC). **(A)** The relationship between SC and age. The vertical axis of the figure represents the proportion of SC connected to each network relative to the whole brain, while the horizontal axis displays age. **(B)** The relationship between FC and age. The vertical axis of the figure represents the mean value of correlation between each network and other region, while the horizontal axis displays age. The regression line on the figure was constructed based on the observed data, and its slope is related to the correlation coefficient (r). The p-value indicates the significance of the relationship between the observed data and the line.

### 3.3 Age related changes in ALFF, global correlation, and local correlation

We investigated the ALFF within each RSN, because the local power of the low-frequency band-passed BOLD signal fluctuation reflects the neuronal activity in the resting state (Nakamura, et al. 2020; Tsurugizawa, et al. 2019). Overall, we observed a positive correlation with age in ALFF of all RSNs (r = 0.47 to 0.87), and ALFF in SN was significantly corelated with age (p = 0.011) (Fig 4). The ALFF in the SMN was positively correlated with age (r = 0.74), but the correlation was not significant (p = 0.060).

**Fig 4.**
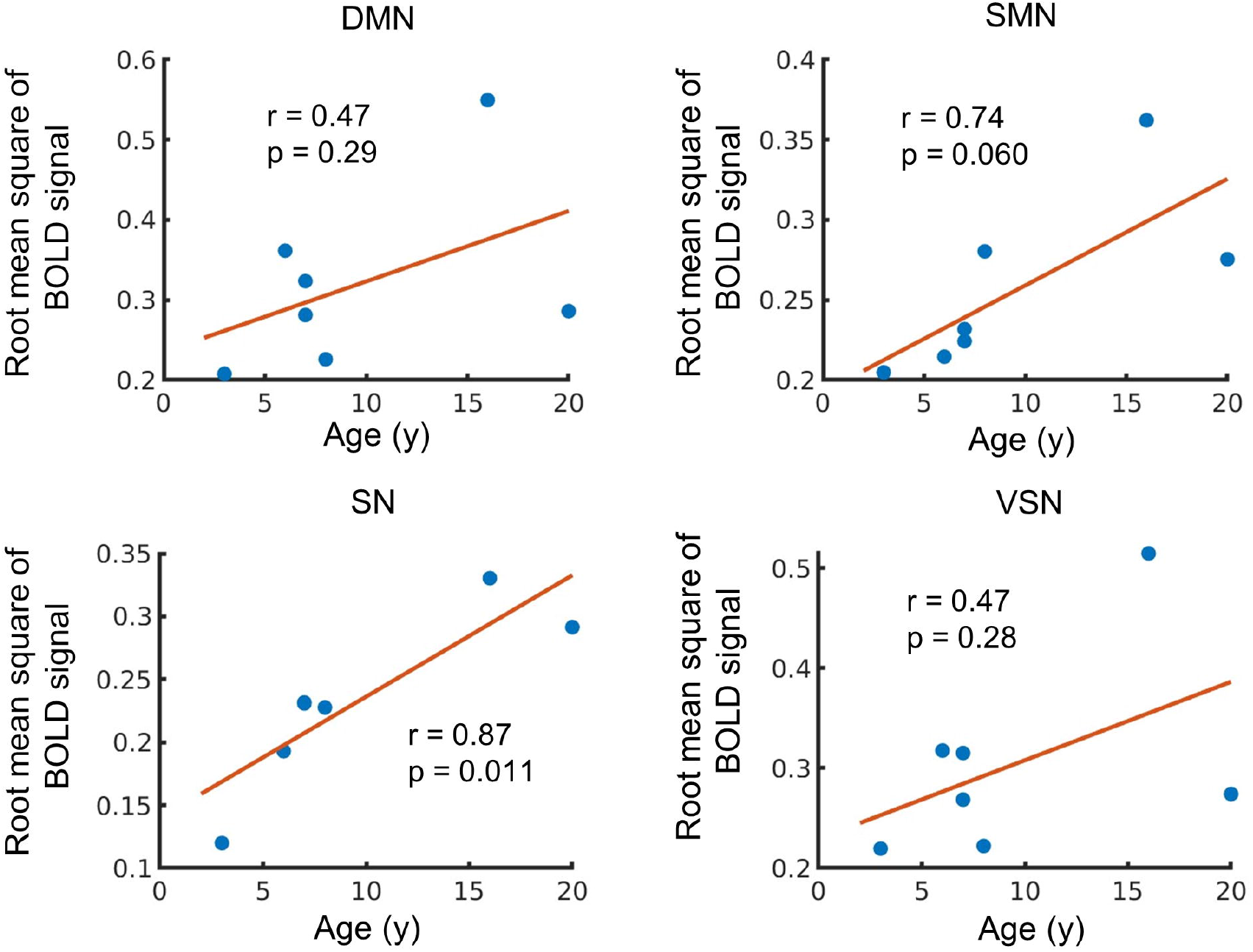
Age-related changes in amplitude low frequency fluctuations (ALFF). The vertical axis of the figure represents the averaged ALFF in each network, while the horizontal axis displays age. The regression line on the figure based on the observed data, and its slope is related to the correlation coefficient (r). The p-value indicates the significance of the relationship between the observed data and the line.

The correlations between the global and local correlations for each RSN were also investigated. A positive correlation between global connectivity and age was observed in the SMN, SN, and VSN, but the correlation was weak and not significant (Fig 5A). A moderate negative correlation in local connectivity in SMN with age was observed, but did not reach significance (r = -0.54 and p = 0.21) (Fig 5B). A weak negative correlation was observed in the DMN, while no correlation was observed in the SN or VSN.

**Fig 5.**
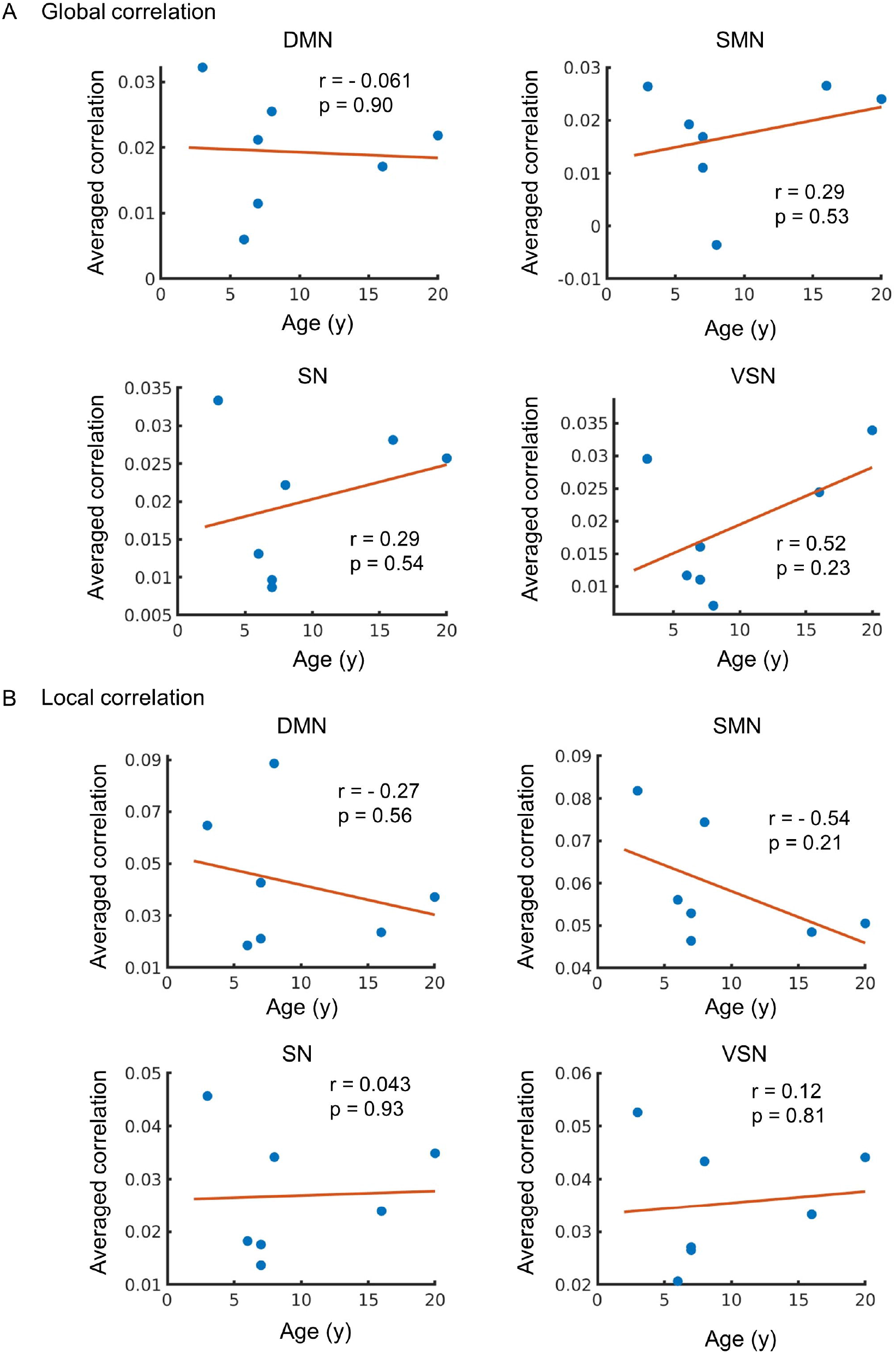
**(A)** The relationship between global correlation and age. The vertical axis of the figure represents the averaged value of global correlation in each network, while the horizontal axis displays age. The regression line on the figure was constructed based on the observed data, and its slope is related to the correlation coefficient (r). **(B)** Age-related changes in local correlation. The relationship between local correlation and age. The vertical axis of the figure represents the averaged value of local correlation, while the horizontal axis displays age. The regression line on the figure based on the observed data, and its slope is related to the correlation coefficient (r). The p-value indicates the significance of the relationship between the observed data and the line.

## 4 Discussion

In the present study, we demonstrated the age-related changes in brain function and structure within the typical RSNs of macaques. A negative correlation between age and SC was identified in specific RSNs, whereas FC and ALFF showed positive correlations with age. As FC and ALFF are caused by neuronal oscillation and activity, these differences could possibly reflect the age-related alterations in the two modalities, such as neuronal activity and brain structure.

### 4.1 Individual similarity of SC and FC

Aged macaques tended to have stronger local connections in the SC than young macaques (Fig 1). The similarity of the SC among macaques aged 5 to 9 years was high (r = 0.87-0.96) (Ouchi, et al. 2024), and the results of this study showed similar findings among young macaques (r = 0.91-0.94). In contrast, the similarity between young and old macaques was relatively low (r = 0.77-0.82) (Fig 1C). Previous studies, which reported that the integrity of the SC decreases with age (Gong et al., 2009), may explain why the similarity in SC among young and old macaques was low (r = 0.77-0.82). Interestingly, the SC between the two old macaques showed higher similarity (r = 0.85) than that between old and young macaques, indicating that structural similarity is still conserved at each life stage.

The overall similarity in FC among young macaques and between young and old macaques was low (r = 0.11-0.25), possibly because anesthesia affects FC (Barttfeld, et al. 2015; Hutchison, et al. 2015; Tsurugizawa and Yoshimaru 2021; Xu, et al. 2019) but not SC. Importantly, although the FC strength is affected by anesthesia, some RSNs are preserved (Tsurugizawa and Yoshimaru 2021). We successfully achieved typical RSNs, including the DMN, SMN, SN, and VSN, under anesthesia, consistent with a previous study (Rao, et al. 2021). Regarding the similarity between old macaques, one prior study hypothesized that age-related reductions in integrity of the SC decrease variation in the FC, indicating that resting-state brain activity may depend on specific connections (Zhang, et al. 2024). The higher similarity between old macaques (r = 0.35) than between young and old macaques (r = 0.11-0.25) may therefore be due to the reduction in the integrity of the SC. A previous study in humans also revealed that the global integration of the FC increases with age (Damoiseaux 2017). However, the present study only observed similarities between the two old macaques. Further investigation using a larger sample size of older macaques is required for further analysis in future studies, although it is difficult to obtain aged macaques.

### 4.2 Age-related changes in RSNs

Here, we focused on four typical networks, namely the DMN, SMN, SN, and VSN, as these RSNs reflect the basal brain networks in the resting state, including in humans. The proportion of SC connected to the DMN, SMN, and VSN in the whole brain decreased with age; however, the percentage of SCs connected to the SN did not change (Fig 3A). Most SCs, including those in the hub regions of the DMN, decline in older subjects in humans (Zimmermann, et al. 2016). The DMN is characterized as being active during wakeful rest compared to when performing a cognitive task in humans and macaques. A previous human study showed that the SC in the SN is less affected by aging (Damoiseaux 2017). The ventral SN is related to affective experiences, whereas the dorsal SN is associated with attention and processing speed (Touroutoglou, et al. 2012). Another study reported that the ventral SN is homologous between humans and macaques (Touroutoglou, et al. 2016). These results are consistent with those of the current study.

In contrast to SC, FC was positively correlated with all RSNs (Fig 3B). A previous study in humans indicated that FC with the DMN in old subjects, defined as those aged 60-83 years old, was greater than that in younger individuals (Grady, et al. 2016; Spreng, et al. 2016). Another study that investigated FC with more classified ages revealed that alterations in FC are not proportional to age. The FC strength and density in the DMN and SMN in over 80 years old group were greater than those in individuals of less than 60 years (Farras-Permanyer, et al. 2019). These results consistent with current study. However, indicate that disparities in the definition of “old” is not strictly decided and the definition of “old” macaques may affect the results of altered FC. This study therefore investigated the correlation of FC with age instead comparing between “young” and “old”, as in previous studies. The present study investigated macaques aging until 20 years old and it is interesting to investigate the brain structure and function in older macaques in the future study.

We further observed a positive correlation between the ALFF and age across all RSNs, indicating an increase in the power of the low-frequency components of the MRI signal in old macaques. Since ALFF is considered a biomarker reflecting neural activity in the resting state (Nakamura, et al. 2020; Tsurugizawa, et al. 2019; Yang, et al. 2007), the positive correlation of ALFF within the SMN and SN with age indicates higher resting-state neural activity in older macaques than in young macaques. A previous human study reported that ALFF increases with age in humans (Zhang, et al. 2021). Furthermore, an increase in the ALFF is correlated with delayed attention/processing speed in the left cerebellar posterior lobe and left superior parietal gyrus, providing evidence that longitudinal cognitive decline shows a frequency-specific correlation with ALFF changes (Fan, et al. 2022). Another study reported that subjects who displayed either low or very high firing rates in layer 2/3 pyramidal cells of the prefrontal cortex exhibited poor performance, whereas those who displayed intermediate firing rates exhibited relatively good performance, indicating that an increase in the action potential firing rate may be responsible, in part, for age-related prefrontal cortex dysfunction (Chang, et al. 2005). As such, the age-related changes in ALFF observed in this study suggest that they may be associated with cognitive decline in macaques. However, it has been reported that anesthesia increases the ALFF in the bilateral prefrontal cortex of macaques (Lv, et al. 2016), and the influence of physiological changes such as respiration and heart rate need to be considered.

Prior studies have shown that local correlation decreases with age and is correlated with language abilities in humans (Pistono, et al. 2021). Additionally, in motor networks, performance has been reported to correlate with local correlation (Wu, et al. 2007). Thus, the results of this study showing that the local correlation in the DMN and SMN decreased with age (Fig 5) may indicate a factor explaining behavioral differences, although it was not statistically significant. In contrast, no correlation with age was observed in the SN or VSN, suggesting that the degree of age-related impact may vary depending on the network. Graph theory studies have demonstrated that local efficiency in resting-state brain activity measured using fMRI decreases with age, whereas global efficiency does not change significantly (Cao, et al. 2014; Geerligs, et al. 2015). Furthermore, local correlations have been reported to explain glucose metabolism in the brain, which decreases with age (Bernier, et al. 2017; Subtirelu, et al. 2023). The present study showed that SC and local correlations within the SMN were negatively correlated with age (Figs. 3-5), which is in agreement with the results of a previous study that showed decreased white matter integrity (Gong, et al. 2009)and glucose metabolic rate (Subtirelu, et al. 2023) in SMN with aging. In contrast, there is a positive correlation between FC and ALFF in all RSNs with age-compensatory spontaneous neural activity increases (Spooner, et al. 2019).

## 5 Conclusion

Typical RSNs such as the DMN of non-human primates are homologous to those in humans; however, MRI study on the age-related changes in FC and SC in macaques are much less common than human studies. The current study showed similar age-related changes in the structure and resting-state function of typical RSNs between humans and macaques, indicating the possibility of performing studies of aging in macaques.

## 7 Author Contributions

**Kazuya Ouchi**: Data Curation, Formal analysis, Investigation, Methodology, Validation, Visualization, Writing - Original Draft, Writing – review and editing. **Shinya Yamamoto**: Investigation, Resources, Funding acquisition, Writing – review and editing. **Makoto Obara:** Resources, Writing – review and editing, **Yasuko Sugase-Miyamoto**: Investigation, Resources, Funding acquisition, Writing – review and editing, **Tomokazu Tsurugizawa**: Conceptualization, Data Curation, Formal analysis, Funding acquisition, Investigation, Methodology, Project administration, Resources, Supervision, Validation, Visualization, Writing – original draft, Writing - Review and Editing.

## 9 Acknowledgments

We thank Mr. Toshiharu Takasu, Mr. Aki Miyamoto, and Ms. Ai Muramatsu for assistance of MRI experiment and taking care of macaques.

## 11 Data Availability Statement

The datasets generated during and/or analyzed during the current study are available from the corresponding author on reasonable request.

## References

Barttfeld, P., et al. 2015 Signature of consciousness in the dynamics of resting-state brain activity. Proc Natl Acad Sci U S A 112(3):887–92.doi:10.1073/pnas.1418031112.

Bernier, M., et al. 2017 Spatial distribution of resting-state BOLD regional homogeneity as a predictor of brain glucose uptake: A study in healthy aging. Neuroimage 150:14–22.doi:10.1016/j.neuroimage.2017.01.055.

Cao, M., et al. 2014 Topological organization of the human brain functional connectome across the lifespan. Dev Cogn Neurosci 7:76–93.doi:10.1016/j.dcn.2013.11.004.

Chang, Y. M., et al. 2005 Increased action potential firing rates of layer 2/3 pyramidal cells in the prefrontal cortex are significantly related to cognitive performance in aged monkeys. Cereb Cortex 15(4):409–18.doi:10.1093/cercor/bhh144.

Damoiseaux, J. S. 2017 Effects of aging on functional and structural brain connectivity. Neuroimage 160:32–40.doi:10.1016/j.neuroimage.2017.01.077.

Dash, S., et al. 2023 Brain volumetrics across the lifespan of the rhesus macaque. Neurobiol Aging 126:34–43.doi:10.1016/j.neurobiolaging.2023.02.002.

Deshpande, G., et al. 2009 Integrated local correlation: a new measure of local coherence in fMRI data. Hum Brain Mapp 30(1):13–23.doi:10.1002/hbm.20482.

Eberling, J. L., et al. 1997 Cerebral glucose metabolism and memory in aged rhesus macaques. Neurobiol Aging 18(4):437–43.doi:10.1016/s0197-4580(97)00040-7.

Fan, D., et al. 2022 Cognitive decline is associated with frequency-specific resting state functional changes in normal aging. Brain Imaging Behav 16(5):2120–2132.doi:10.1007/s11682-022-00682-1.

Farras-Permanyer, L., et al. 2019 Age-related changes in resting-state functional connectivity in older adults. Neural Regen Res 14(9):1544–1555.doi:10.4103/1673-5374.255976.

Geerligs, L., et al. 2015 A Brain-Wide Study of Age-Related Changes in Functional Connectivity. Cereb Cortex 25(7):1987–99.doi:10.1093/cercor/bhu012.

Goelman, G., et al. 2024 Directed functional connectivity of the default-mode-network of young and older healthy subjects. Sci Rep 14(1):4304.doi:10.1038/s41598-024-54802-6.

Gong, G., et al. 2009 Age- and gender-related differences in the cortical anatomical network. J Neurosci 29(50):15684–93.doi:10.1523/JNEUROSCI.2308-09.2009.

Grady, C., et al. 2016 Age differences in the functional interactions among the default, frontoparietal control, and dorsal attention networks. Neurobiol Aging 41:159–172.doi:10.1016/j.neurobiolaging.2016.02.020.

Hutchison, R. M., et al. 2015 Functional subdivisions of medial parieto-occipital cortex in humans and nonhuman primates using resting-state fMRI. Neuroimage 116:10–29.doi:10.1016/j.neuroimage.2015.04.068.

Hutchison, R. M., et al. 2011 Resting-state networks in the macaque at 7 T. Neuroimage 56(3):1546–55.doi:10.1016/j.neuroimage.2011.02.063.

Hyon, R., et al. 2020 Similarity in functional brain connectivity at rest predicts interpersonal closeness in the social network of an entire village. Proc Natl Acad Sci U S A 117(52):33149–33160.doi:10.1073/pnas.2013606117.

Lv, P., et al. 2016 Dose-dependent effects of isoflurane on regional activity and neural network function: A resting-state fMRI study of 14 rhesus monkeys: An observational study. Neurosci Lett 611:116–22.doi:10.1016/j.neulet.2015.11.037.

Nakamura, Y., et al. 2020 fMRI detects bilateral brain network activation following unilateral chemogenetic activation of direct striatal projection neurons. Neuroimage 220:117079.doi:10.1016/j.neuroimage.2020.117079.

Noda, A., et al. 2002 Age-related changes in cerebral blood flow and glucose metabolism in conscious rhesus monkeys. Brain Res 936(1-2):76–81.doi:10.1016/s0006-8993(02)02558-1.

Ouchi, Kazuya, et al. 2024 Multi-scale hierarchical brain regions detect individual and inter-species variations of structural connectivity in macaque monkeys and humans. PREPRINT at Research Square.doi:10.21203/rs.3.rs-4092810/v1.

Pistono, A., et al. 2021 Increased functional connectivity supports language performance in healthy aging despite gray matter loss. Neurobiol Aging 98:52–62.doi:10.1016/j.neurobiolaging.2020.09.015.

Rao, B., et al. 2021 Development of functional connectivity within and among the resting-state networks in anesthetized rhesus monkeys. Neuroimage 242:118473.doi:10.1016/j.neuroimage.2021.118473.

Saad, Z. S., et al. 2013 Correcting brain-wide correlation differences in resting-state FMRI. Brain Connect 3(4):339–52.doi:10.1089/brain.2013.0156.

Smith, S. M., et al. 2004 Advances in functional and structural MR image analysis and implementation as FSL. Neuroimage 23 Suppl 1:S208–19.doi:10.1016/j.neuroimage.2004.07.051.

Spooner, R. K., et al. 2019 Rhythmic Spontaneous Activity Mediates the Age-Related Decline in Somatosensory Function. Cereb Cortex 29(2):680–688.doi:10.1093/cercor/bhx349.

Spreng, R. N., et al. 2016 Attenuated anticorrelation between the default and dorsal attention networks with aging: evidence from task and rest. Neurobiol Aging 45:149–160.doi:10.1016/j.neurobiolaging.2016.05.020.

Stumme, J., et al. 2022 Interrelating differences in structural and functional connectivity in the older adult’s brain. Hum Brain Mapp 43(18):5543–5561.doi:10.1002/hbm.26030.

Subtirelu, R. C., et al. 2023 Aging and Cerebral Glucose Metabolism: (18)F-FDG-PET/CT Reveals Distinct Global and Regional Metabolic Changes in Healthy Patients. Life (Basel) 13(10).doi:10.3390/life13102044.

Tomasi, D., G. J. Wang, and N. D. Volkow 2013 Energetic cost of brain functional connectivity. Proc Natl Acad Sci U S A 110(33):13642–7.doi:10.1073/pnas.1303346110.

Touroutoglou, A., et al. 2016 A ventral salience network in the macaque brain. Neuroimage 132:190–197.doi:10.1016/j.neuroimage.2016.02.029.

Touroutoglou, A., et al. 2012 Dissociable large-scale networks anchored in the right anterior insula subserve affective experience and attention. Neuroimage 60(4):1947–58.doi:10.1016/j.neuroimage.2012.02.012.

Tsurugizawa, T. B. Djemai, and A. Zalesky 2019 The impact of fasting on resting state brain networks in mice. Sci Rep 9(1):2976.doi:10.1038/s41598-019-39851-6.

Tsurugizawa, T., and D. Yoshimaru 2021 Impact of anesthesia on static and dynamic functional connectivity in mice. Neuroimage 241:118413.doi:10.1016/j.neuroimage.2021.118413.

Upright, N. A., and M. G. Baxter 2021 Prefrontal cortex and cognitive aging in macaque monkeys. Am J Primatol 83(11):e23250.doi:10.1002/ajp.23250.

Whitfield-Gabrieli, S., and A. Nieto-Castanon 2012 Conn: a functional connectivity toolbox for correlated and anticorrelated brain networks. Brain Connect 2(3):125–41.doi:10.1089/brain.2012.0073.

Wu, T., et al. 2007 Normal aging decreases regional homogeneity of the motor areas in the resting state. Neurosci Lett 423(3):189–93.doi:10.1016/j.neulet.2007.06.057.

Xu, T., et al. 2019 Interindividual Variability of Functional Connectivity in Awake and Anesthetized Rhesus Macaque Monkeys. Biol Psychiatry Cogn Neurosci Neuroimaging 4(6):543–553.doi:10.1016/j.bpsc.2019.02.005.

Yang, H., et al. 2007 Amplitude of low frequency fluctuation within visual areas revealed by resting-state functional MRI. Neuroimage 36(1):144–52.doi:10.1016/j.neuroimage.2007.01.054.

Zhang, H., X. Bai, and M. T. Diaz 2021 The intensity and connectivity of spontaneous brain activity in a language network relate to aging and language. Neuropsychologia 154:107784.doi:10.1016/j.neuropsychologia.2021.107784.

Zhang, H., et al. 2024 The structural-functional-connectivity coupling of the aging brain. Geroscience 46(4):3875–3887.doi:10.1007/s11357-024-01106-2.

Zimmermann, J., et al. 2016 Structural architecture supports functional organization in the human aging brain at a regionwise and network level. Hum Brain Mapp 37(7):2645–61.doi:10.1002/hbm.23200.

